# Understanding phase-amplitude coupling from bispectral analysis

**DOI:** 10.1101/2020.03.20.000745

**Authors:** Coen S. Zandvoort, Guido Nolte

## Abstract

Two measures of cross-frequency coupling (CFC) are Phase-Amplitude Coupling (PAC) and bicoherence. The estimation of PAC with meaningful bandwidth for the high frequency amplitude is crucial in order to avoid misinterpretations. While recommendations on the bandwidth of PAC’s amplitude component exist, there is no consensus yet. Here, we show that the earlier recommendations on filter settings lead to estimates which are smeared in the frequency domain, which makes it difficult to distinguish higher harmonics from other types of CFC. We also show that smearing can be avoided with a different choice of filter settings by theoretically relating PAC to bicoherence. To illustrate this, PAC estimates of simulations and empirical data are compared to bispectral analyses. We used simulations replicated from an earlier study and empirical data from human electro-encephalography and rat local field potentials. PAC’s amplitude component was estimated using a bandwidth with a ratio of (1) 2:1, (2) 1:1, or (3) 0.5:1 relative to the frequency of the phase component. For both simulated and empirical data, PAC was smeared over a broad frequency range and not present when the estimates comprised a 2:1- and 0.5:1-ratio, respectively. In contrast, the 1:1-ratio accurately avoids smearing and results in clear signals of CFC. Bicoherence estimates, which do not smear across frequencies by construction, were found to be essentially identical to PAC calculated with the recommended frequency setting.

## 1. Introduction

Electroencephalography (EEG), magnetoencephalography (MEG), and local field potentials (LFPs) are often used to measure brain activity. They allow for the registration of neuronal population activity with high temporal precision. Associations within and between neural populations are regularly assessed using second-order measures, like power- and cross-spectral estimates. However, such measures are constrained to merely reveal linear interactions between signals. It does not come as a surprise that brain activity also contains neural dynamics related to nonlinear processes (Florin and Baillet [2015]; Jensen and Colgin [2007]; Tort et al. [2007]). To unravel these dynamics, nonlinear measures have gained much popularity in recent decades. In general, measures of cross-frequency coupling (CFC) describe the nonlinear interactions of signals across frequencies, being it coupling across amplitude, phase, or frequency. A CFC-measure that was rather unexplored but gained increasingly popularity to assess phase-amplitude interactions is bicoherence (Bartz et al. [2019]; Shahbazi-Avarvand et al. [2018]). Bicoherence describes the normalised cross-bispectrum. The cross-bispectrum is a measure of CFC since it evaluates the interactions across three frequencies. The first two frequencies can be chosen at will and the third frequency is always constrained to be the sum of these two frequencies. In statistical terms, the cross-bispectrum is the third-order moment in the frequency domain. Similar to coherence, bicoherence is a measure of phase-phase coupling even though it also somewhat depends on amplitudes because trials with larger amplitudes receive a larger weight in the average. This dependence on amplitudes should be distinguished from the observation by Hyafil [2015] that bicoherence is strongly related to phase-amplitude coupling because specific phase relations can in general generate an amplitude modulation. This is well known in acoustics as the phenomenon of beating where, for example, two sinusoidal sounds with similar frequencies lead to a modulated amplitude with the difference of these frequencies. In this setting, the modulation of one amplitude and the relation between two phases describe the same phenomenon in different terminology. If such an amplitude modulation is related to a phase of a low frequency signal, then, this phenomenon can be described by both a relation of a low frequency phase and a high frequency amplitude and also by the relation of three phases at three different frequencies.

It is somewhat confusing that the term “phase-amplitude coupling” is used interchangeably in the literature to mean a specific phenomenon and also specific measures used to detect this phenomenon. The construction of those measures is guided by the idea of the mechanism: first, amplitudes and phases are extracted from the data, and then the functional relation between these signals is calculated. Here, we show in much detail that both bicoherence and PAC-measures detect PAC as a phenomenon, but we will reserve the abbreviation PAC for those measures which are defined explicitly as a coupling between phase and amplitude.

Although PAC and bicoherence share similarities, it has been argued that bicoherence has several advantages over PAC (Shahbazi-Avarvand et al. [2018]). One advantage of bicoherence over PAC is that it is not necessary to filter the signals before CFC computation, since Fourier transforms are iteratively performed over isolated frequencies. Earlier investigations also emphasized that the filter characteristics (i.e. centre frequencies and bandwidth) of PAC have effects on the subsequent estimates (Aru et al. [2015]; Berman et al. [2012]). Hence, filter features selected for both the low- and high-frequency signals can affect the detection of PAC significantly (Berman et al. [2012]). The bandwidth of the high-frequency component is critical in accurately estimating PAC. A bandwidth that is chosen too narrow or broad is prone to statistical errors (i.e. result in false negatives and positives, respectively). The optimal bandwidth settings remain to be examined further. A recent investigation recommended that the bandwidth of the amplitude component should be at least twice as large as the frequency of the phase component (Aru et al. [2015]). This recommendation was followed by experimental studies (Cheng et al. [2016]; Martínez-Cancino et al. [2019]; Murta et al. [2017]; Seymour et al. [2017]). If one aims to investigate narrow-band brain rhythms and its higher harmonics, however, a bandwidth that is twice as large as the phase component would result in the inclusion of multiple carrier frequencies. Although this would result in significant PAC, clear smearing across neighbouring frequencies will be shown. To find associations in brain rhythms that are operating within narrow frequency bands would therefore be too complicated for PAC. A typical example of a narrow-band rhythm is the alpha rhythm (8-12 Hz) and its higher harmonics in closed-eyes human resting state. PAC may prevent us to find coupling in such rhythms with a spectrally narrowed operational window.

In the current study, we show that the claim of taking a bandwidth of PAC’s amplitude component that is twice as large as the frequency of the phase component is neither necessary nor optimal. To prove this, we focus on phase-amplitude coupling from a bispectral analysis point of view, and compare bicoherence and PAC analytically and findings of both simulations and empirical data. First, we shortly recall how one can understand PAC using bicoherence, after which we replicate the simulations performed in the earlier investigation for both PAC and bicoherence (Aru et al. [2015]). Specifically, we show how bicoherence relates to PAC filtered with varying bandwidths for the amplitude component. PAC’s components are filtered in a ratio of 2:1, 1:1, and 0.5:1 between the amplitude component and frequency of the phase component. Empirical findings using human EEG and rat LFPs are reported to substantiate the simulation findings. For real EEG and LFPs data, we demonstrate how results for PAC depend more heavily on these ratios when one is interested in narrow-band neural dynamics, and how that can be understood from bicoherence analysis.

## 2. Methods

### 2.1. Phase-amplitude coupling and bicoherence definitions

#### 2.1.1. Phase-amplitude coupling

Several versions of PAC exist, and we first recall the formal definition of PAC given in Canolty et al. [2006]. The essential quantity for PAC, which we consider as a weighted phase locking, is

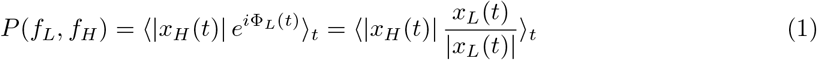

where *x_L_*(*t*) and *x_H_*(*t*) are the (complex) Hilbert transforms of a signal filtered in a low and high frequency band with centers at frequencies *f_L_* and *f_H_*, respectively. We here defined *P* as a complex number and consider its absolute value as a coupling measure if needed. The corresponding phases and magnitudes at time t are denoted by Φ*_L/H_*(*t*) and |*x_L/H_*(*t*)|, and 〈·〉_*t*_ denotes the average over time. From *P* and corresponding values of that quantity for surrogate data, *P_s_*, where the low and a high frequency parts are shifted relative to each other by a random delay, Canolty et al. [2006] calculate a z-score as

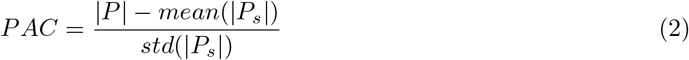

where *mean*(|*P_s_*|) and *std*(|*P_s_*|) are the mean and standard deviations of |*P_s_*| calculated of *K* surrogate data sets. In our opinion, the calculation of this z-score and its interpretation as a Gaussian distributed variable with zero mean and unit standard deviation is highly problematic, because |*P*| is not Gaussian distributed. An alternative statistics will be discussed in section 2.5. Apart from details how significance is estimated, the crucial point is that in this approach PAC is normalized statistically and results depend systematically on the number of trials.

A different approach was suggested by Osipova et al. [2008] where the coherence between the instantaneous high frequency power and the original data was calculated at some low frequency. A difference is that coherence includes a normalization while Canolty et al. [2006] only use a statistical normalization, i.e. coupling measures are normalized by estimates of the standard deviation. Here, we will present results using both a conventional and statistical normalization, and we therefore essentially adopted the approach by Osipova et al. [2008] with an additional statistical normalization.

Another difference is that instantaneous power is calculated by Osipova et al. [2008] with a wavelet approach rather than Hilbert-transformed high frequency filtered data. We slightly deviate from the approach by Osipova et al. [2008] by using high frequency amplitude rather than power and we used the absolute value of the Hilbert transform of the filtered data rather than wavelets to estimate the time-dependent amplitude, because we then have more freedom to design an appropriate filter. In particular, we used a filter which is essentially flat within the high frequency band, whereas the wavelet approach corresponds to a filter which is maximal in the centre of the band and decreases with distance from the centre.

Relations for PAC and bispectra were discussed in [Shahbazi-Avarvand et al., 2018], and we want to first show that the approach of Osipova et al. [2008] leads to the same relation. We start with relating the wavelet approach with the approach based on the Hilbert transform. In a wavelet approach, the complex signal at frequency *f_H_* is calculated as

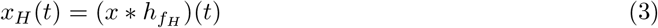

where * denotes convolution and *h_f_H__*(*t*) is the wavelet.

To simplify the notation, we consider an odd number of discrete time points running from −*N* to *N*, and the Fourier transforms, to be used below, are defined for discrete frequencies *f* also running from −*N* to *N*. With such a convention the wavelet is defined as

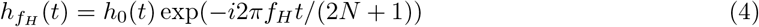

with *h*_0_(*t*) chosen by Osipova et al. [2008] to be a Hanning window. PAC is finally defined in that approach as the coherence at some low frequency *f_L_* between the original data and the instantaneous power

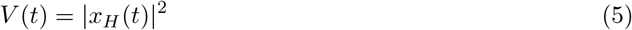

In the context of PAC we prefer the Hilbert approach over the wavelet approach, but there is no fundamental difference between them. To see this, let 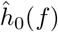 be the Fourier transform of the window, which is substantially different from zero only for small frequencies. Then the Fourier transform of the wavelet reads

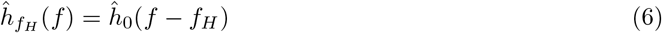

The convolution in Eq.3 is a product in the Fourier domain, and hence

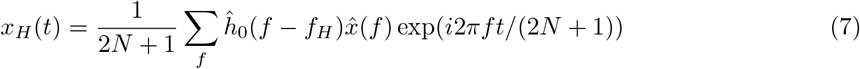

The crucial point now is that, while 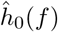 does not vanish for negative frequencies, 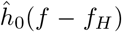 is negligible for negative frequencies provided that the high frequency *f_H_* is remote from zero and the Nyquist frequency relative to the width of the wavelet in the frequency domain. We here assume that this is an excellent approximation in practice, and using this approximation we can write

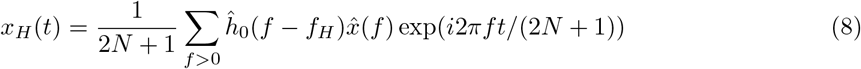

which only differs from Eq.7 by the range of the sum over *f*. This has the form of a Hilbert transform of a filtered signal with the filter given by 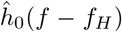. Thus, the wavelet approach is, to excellent approximation, equivalent to the approach based on the Hilbert transformation. This result is almost identical to the main result of Bruns [2004], where it is argued that these two approaches are formally equivalent. The only difference is that we do not think that the derivation given by Bruns [2004] is accurate, but contains an approximation at some point, which should be mentioned even if the approximation is usually very good.

#### 2.1.2. Bispectra and bicoherence

The cross-bispectrum is a poly-spectra, or more specifically, the third-order statistical moment of the Fourier transform. The cross-bispectrum reads in the most general form for channels *k*, *m*, *n* and frequencies *f*_1_ and *f*_2_:

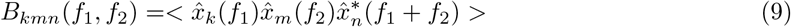

where 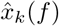 denotes the Fourier transform of data in channel *k* and frequency *f*, and <> denotes the expectation value, which can, for example, be estimated using a windowed segmentation approach.

The cross-spectrum is a nonlinear measure of a single time series across multiple frequencies. To be able to interpret bicoherence like coherence, the cross-bispectrum is here normalised with a three-norm defined as

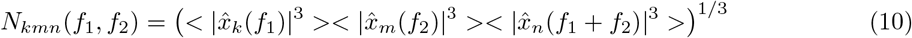

to obtain a coupling measure with absolute value bounded between 0 and 1 (Shahbazi-Avarvand et al. [2014]). Bicoherence is obtained by dividing the cross-bispectrum of Eq.9 by normalisation as described in the equation above:

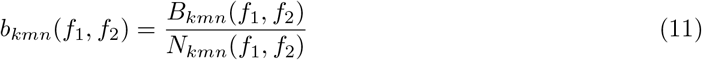

The code to estimate univariate bicoherence is open-source available in the MATLAB-based METH-toolbox, which can be downloaded from the website of the Department of Neurophysiology and Pathophysiology at the Univerisitätklinikum Hamburg-Eppendorf (https://www.uke.de/english/departments-institutes/institutes/neurophysiology-and-pathophysiology/research/research-groups/index.html).

### 2.2. Relations between Phase-Amplitude Coupling and bispectra

#### 2.2.1 PAC as amplitude weighted phase locking

Relations between PAC and bispectra will be formulated for the univariate case. The multivariate case just corresponds to renaming variables and will also be discussed below. To find exact relationships with bispectra it is necessary to slightly redefine PAC such that it can be expressed in terms of third order statistical moments. In particular, the amplitude needs to be replaced by the power (the square of the amplitude) and the low frequency phase needs to be weighted by the low frequency amplitude. A corresponding modification of the measure defined in Eq.1 reads

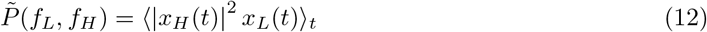

*F_L_*(*f*, *f_L_*) and *F_L_*(*f*, *f_H_*) can be denoted as the filters for the low and high frequency part as a function of frequency *f* and with centers *f_L_* and *f_H_*. With

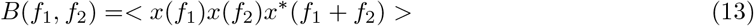

being the bispectrum at frequencies *f*_1_ and *f*_2_ it was shown in [Shahbazi-Avarvand et al., 2018] that it relates to 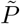 as

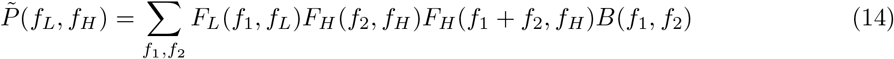

apart from an irrelevant overall multiplicative constant.

While the width of the filter for the high frequency part is much debated, the filter for the low frequency part can in principle be arbitrarily narrow and is only limited by the trade-off between frequency resolution and statistical robustness. To discuss the implications of the above relation we will therefore introduce a simplification and assume that the low frequency filter is point-like in the frequency domain, i.e.

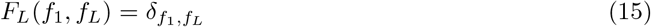

which leads to

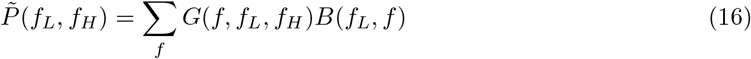

with the kernel

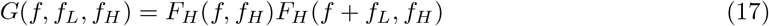

The kernel is a product of the high frequency filter and a shifted version of it, and hence, 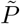 is in general a smeared version of *B*.

#### 2.2.2 PAC as coherence between instantaneous power and original data

In Osipova et al. [2008], PAC was defined as coherence between original data and instantaneous paper. Similar to the preceding section, relations between PAC and bispectra can be given only for the numerator and not for the normalization factors. Also, we here define the instantaneous power using the Hilbert transform rather than using wavelets.

The Hilbert transform of filtered data in the high frequency band reads

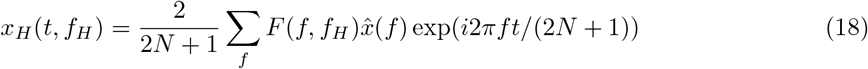

where *F*(*f*, *f_H_*) is the filter with center *f_H_* which is assumed here to be real valued. Also, in this formulation *F*(*f*, *f_H_*) is set to zero for negative *f*. Using the orthogonality of sinusoidal functions

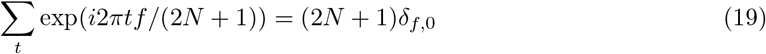

then the Fourier transform of the power *V*(*t*, *f_H_*) ≡ |*x_H_*(*t*, *f_H_*)|^2^ at some frequency *f_L_* reads

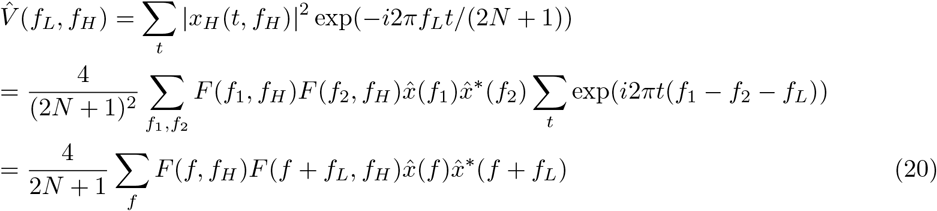

The cross-spectrum, *c*(*f_L_*, *f_H_*), at frequency *f_L_* between the original data and the power now reads

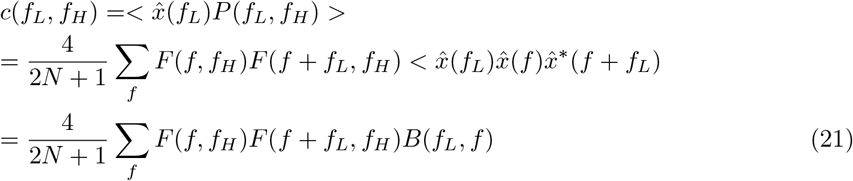

which is identical to Eq.16 if we ignore the irrelevant constant 4/(2*N* +1).

#### 2.2.3 Setting the high frequency filter

The smearing is negligible if the kernel *G* in Eq.17 is approximately equal to a delta-function. For perfect high frequency filters, i.e. if the filters are 1 within some band and vanish outside that band, the kernel is approximately equal to a delta-function if the width is chosen to be slightly larger than the low frequency *f_L_*. The filter and its shifted version then only overlap in a very narrow frequency region and the kernel has a sharp peak. More formally, if

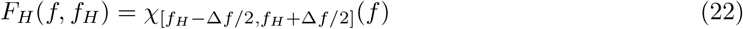

with Δ*f* being the bandwidth and 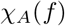 being the characteristic function on the interval *A*, and choosing Δ*f* = *f_L_* + *ϵ* with a tiny number *ϵ*, then

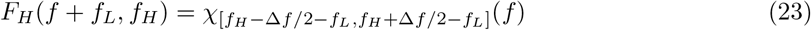

and

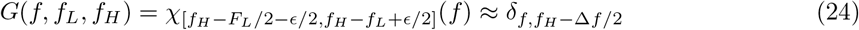

leading to

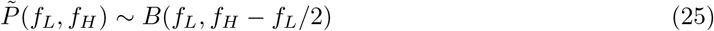

Note that in this case 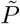 and *B* are identical apart from a frequency shift by *f_L_*/2 and apart from an overall constant, which depends on the filter and is usually irrelevant. The frequency shift is due to different conventions regarding what is meant by a frequency. That shift would disappear if PAC would be written as a function of the lower end of the filter rather than the center. For example, the first higher harmonic of the alpha rhythm (i.e. the coupling between 10 Hz and 20 Hz), could be observed with PAC with a bandwidth of 10 Hz for the high frequency filter at low frequency *f_L_* = 10 Hz and high frequency *f_H_* = 15 Hz, because then the high frequency band includes both signals at 10 and 20 Hz. In contrast, the same phenomenon can be observed with the bispectrum at *f*_1_ = *f*_2_ = 10 Hz, which corresponds to a coupling between 10 Hz and *f*_1_ + *f*_2_ = 20 Hz. Note that displaying PAC as a function of the center of the high frequency band can be misleading in the case of higher harmonics because we would observe a peak at 15 Hz even though there is no relevant signal at 15 Hz.

In practice, filters are not perfect, and it is not necessary to choose the filter width slightly larger than the low frequency. Rather, choosing the width of the high frequency filter to be equal to the low frequency results in sharp kernels (with details depending on filter order) and avoids smearing of frequencies. The effect of the filter width on the kernel *G* is shown in Fig.1. This is the main theoretical result of this paper which will be illustrated below for simulated and empirical data. In particular, Kovach et al. [2018] derived relations between bispectra and (unnormalized) PAC and argued that the quantities are related but not identical. We here disagree with the latter as a very general statement: in principle, with a proper choice of filters the quantities are identical. We finally recall that such relations are only valid for the numerators of the coupling measures, but for bicoherence it was shown in Shahbazi-Avarvand et al. [2018] and Shahbazi-Avarvand et al. [2014] that different normalizations have almost no effect on its statistical properties.

**Figure 1:**
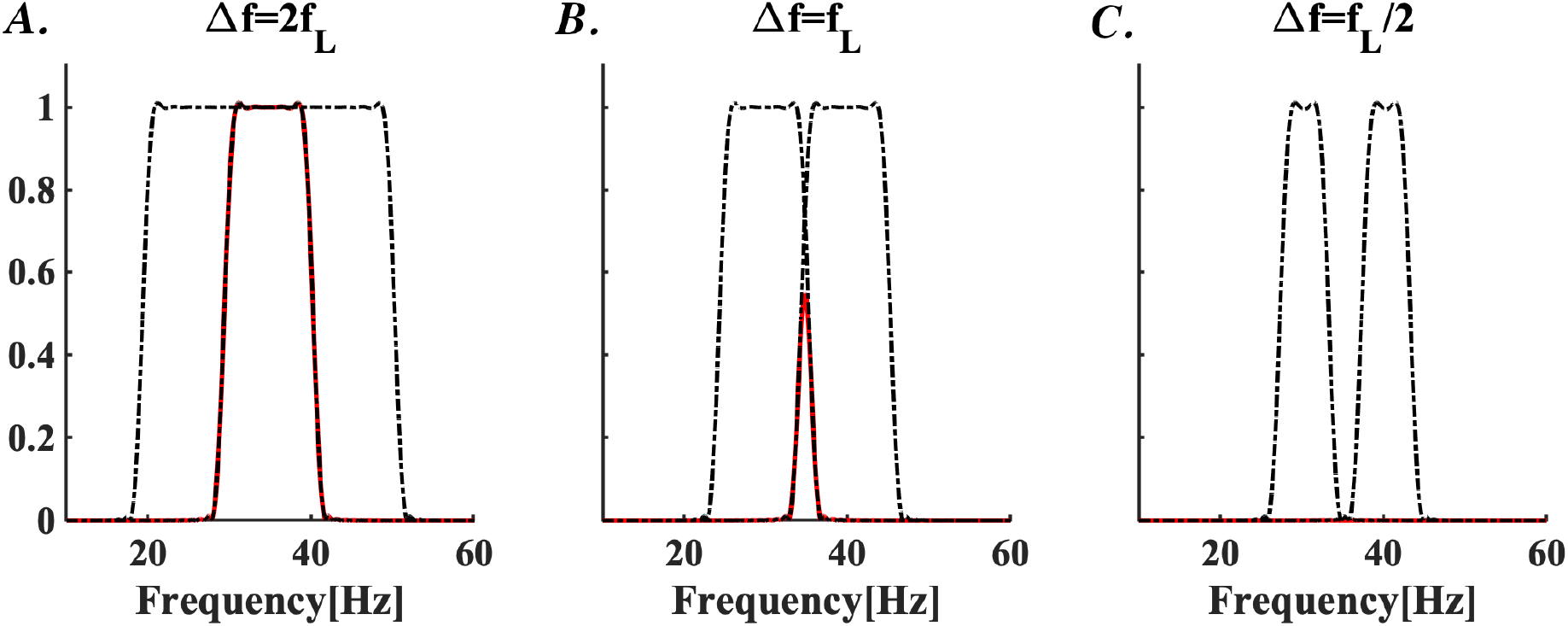
Filters were calculated for fixed low frequency *f_L_* = 10 Hz and high frequency *f_H_* = 40 Hz for varying width of the high frequency band, denoted as Δ*f* and indicated in the titles of the subpanels. The filters and the corresponding shifted filters are shown by black lines. The kernels *G*, i.e. the products of the filters, are shown in red. We used a FIR filter of 1 second duration. At B) Δ*f* = *f_L_* we observe a (moderately) sharp kernel. For A) Δ*f* = 2*f_L_* the kernel is not sharp, and for C) Δ*f* = *f_L_*/2 it basically vanishes altogether. Of course, the kernel can be made sharper with longer filters.

As the primary aim is to reveal the importance of correctly selecting an adequate filter width for PAC, PAC’s amplitude component will be filtered below with three bandwidths. First, the recommendation of Aru and colleagues is followed. Recall that they recommend to filter the amplitude component with a bandwidth that is twice as large as the frequency of the phase component. Second, as spectral leakage (i.e. smearing) over a broad frequency band is expected for narrow-band coupling, we selected the amplitude bandwidth to be identical to the frequency of the phase component. Finally, to show that the bandwidth cannot be too small, we additionally filtered the amplitude component with a bandwidth that is half the frequency of the phase component. In other words, the amplitude component of PAC is filtered in a 2:1-, 1:1-, and 0.5:1-ratio to the frequency of the phase component. For simplicity, we only focus on the univariate cases of bicoherence and PAC.

#### 2.2.4 Remarks on the multivariate case

The theory presented above is completely analogous for the bivariate case. If the low frequency signal is taken from channel *k*, and the high frequency signal from channel *m*, then it is straightforward to show that Eq.25 changes to

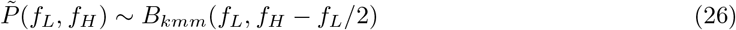

where we only added channel indices in the bispectra. This shows that PAC is a special case of bispectra: while PAC is a bivariate measure, bispectra are, in general, trivariate measures where three channel indices can be set freely. Additionally, it was shown in Chella et al. [2014] that the antisymmetric combination *B_kmm_* – *B_mkm_* is robust to artefacts of volume conduction (i.e. a significantly non-vanishing result on channel level is inconsistent with a mixing of independent sources). Such a quantity cannot be calculated with PAC because no corresponding term for *B_mkm_* exists.

### 2.3. Simulated data

We first focus on the simulations as performed by Aru and colleagues. In their study, they simulated a univariate signal oscillating at different frequencies. More precisely, the simulated signals manifested PAC dynamics because the amplitude of the high frequency signal was modulated by the phase of the low frequency signal. We formulate the following time series which is analogous to theirs:

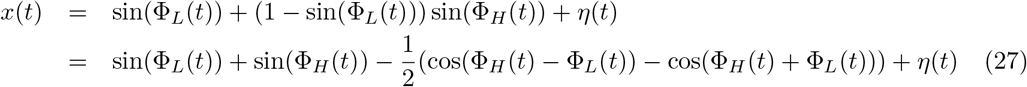

where Φ_*L*_(*t*) and Φ_*H*_(*t*) denote the phase time series, and *η*(*t*) is white Gaussian noise with a standard deviation of 3. The noise is added to avoid oscillating effects at irrelevant frequencies merely due to spectral leakage. We note that taking the phases as Φ_*L,H*_(*t*) = 2*πf_L,H_t* would have resulted in comparable results in our case. However, taking pure sine waves in simulations should be avoided because of principal reasons. Pure sine waves have infinite auto-correlation length and, hence, if data are divided into epochs, all epochs are statistically dependent on each other. In fact, pure sine waves are not ergodic, meaning that the time averages are generally not identical to ensemble averages. As an example, consider two perfect sine waves of identical frequency but random and independent initial phases. The coherence between the two sine waves is 1 when calculated as a time average, because the phase difference is constant across time whereas it vanishes when calculated as an ensemble average, because the phases are random. Here, the phase time series are obtained from the phase of the Hilbert transforms of white Gaussian noise which was narrowly filtered around the frequencies of 10 and 40 Hz. We note that the simulated time series contain additional dominant frequencies at 30 and 50 Hz because of the modulated high-frequency component.

### 2.4. Empirical electro-encephalographic and local field potential data

To substantiate our simulation findings, we compared estimates of bicoherence and PAC on empirical data recorded in humans and rats. Both datasets are open-source and publicly available on http://clopinet.com/causality/data/nolte/ and doi.org/10.5061/dryad.12t21. The human data comprise cleaned resting state EEG. The online dataset is a subset of data that was reported earlier (Nolte et al. [2008]), since the original dataset consisted of 88 participants. In the experimental paradigm, participants were asked to close their eyes for 10-13 minutes. Simultaneously, EEG recordings were acquired, which were epoched into 152-202 segments of 4s each. 19-channel EEG (configured to the international 10-20 system) was registered at a sampling rate of 256 samples/s. EEG data are referenced to a linked mastoids reference. Nineteen Ag/AgCl electrodes included Fp1, Fp2, F7, F3, Fz, F4, F8 T3, C3, Cz, C4, T4, T5, P3, Pz, P4, T6, O1, and O2. Given that we were interested in the coupling of narrow-band brain rhythms, the primary aim was to localise neural oscillations at the alpha rhythm and its higher harmonics. To find the strongest coupling effects, we specifically targeted the EEG channel comprising maximal alpha power (around 10 Hz). We selected channel 17 of participant 8 (i.e. the channel yielding maximal alpha power) for further analysis. The alpha oscillations have a ground frequency around 10 Hz. The higher harmonics yield frequencies of a factor k*10 higher (with k=1,2,3,…). Rat data comprise LFPs of seven animals. LFP data were recorded during 12 hours of sleep and awake periods. We report findings on the REM sleeping periods. LFP activity was recorded from the CA1 pyramidal cell layer of the hippocampus. Data of 16-channel silicon-layered probes were sampled at a rate of 1,000 Hz. In the original investigation, the authors mainly aimed to identify theta-gamma PAC (and cross-frequency phase-phase coupling) in hippocampal activity (Scheffer-Teixeira and Tort [2016]). We predominantly focus on the theta-gamma phase-amplitude coupling, which would be the cross-frequency coupling between the frequencies of 4-8 and 40-150 Hz. For the rat findings, we specifically report the results of channel 16 of rat 7.

### 2.5. Statistical assessments

In order to statistically assess single PAC and bicoherence representations, we applied a surrogate technique. The null-hypothesis is defined as the coupling being zero. More specifically, it is assessed whether non-zero values of PAC and bispectra can be considered as meaningful phase-amplitude coupling or as findings by chance. The surrogate analysis consisted of time-shifting the amplitude component of PAC and shuffling the third Fourier transform of cross-bispectrum. For the construction of the surrogate data sets we follow the approach of Canolty et al. [2006] but deviate in the statistical interpretation of the results. For PAC, we randomly time-shift the amplitude component by a random multiple of 1s with periodic boundary condition, and for the bispectra the same is done for the signal at the highest (i.e. at frequency *f*_1_ + *f*_2_). Using the central limit theorem, under the null-hypothesis *P* is approximately Gaussian distributed in the complex plane with zero mean and uniformly distributed phase. Then |*P*|^2^ is exponentially distributed with density *pdf*(|*P*|^2^) ~ exp(|*P*|^2^/*α*^2^) and with *α*^2^ =< |*P*|^2^ > (Freyer et al. [2009]). Furthermore, under the null-hypothesis *P* is drawn from the same ensemble as the surrogates, and hence *α*^2^ can be estimated as

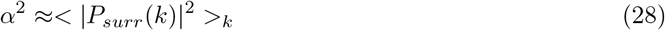

where *P_surr_*(*k*) is the coupling of the *k.th* surrogate data set, and <>_*k*_ denotes average over all surrogate data sets. In the results shown below we will display the quantity

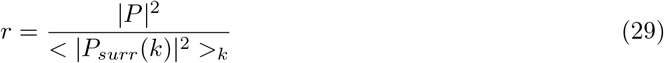

and it is straightforward to show, by integrating the probability density, that the p-values can be calculated as *p* = exp(−*r*). Statistical results for bispectra were calculated analogously. Note, for both PAC and bispectra, we used unnormalized quantities for statistical evaluation because it is these quantities which are plane averages over segments and which are then approximately Gaussian distributed in the complex plane.

## 3. Results

### 3.1. Simulated data

Examples of the simulated data and its power spectral density are depicted in Fig. 2A and B, respectively. The power spectral density highlights peaks at 10, 30, 40, and 50 Hz. Although our simulated data only contained two frequencies (i.e. 10 and 40 Hz), the two side peaks at 30 and 50 Hz were introduced due to the 40-Hz modulation. To be able to identify PAC, at least one of these two side frequencies must be included in the signal of the amplitude component.

**Figure 2:**
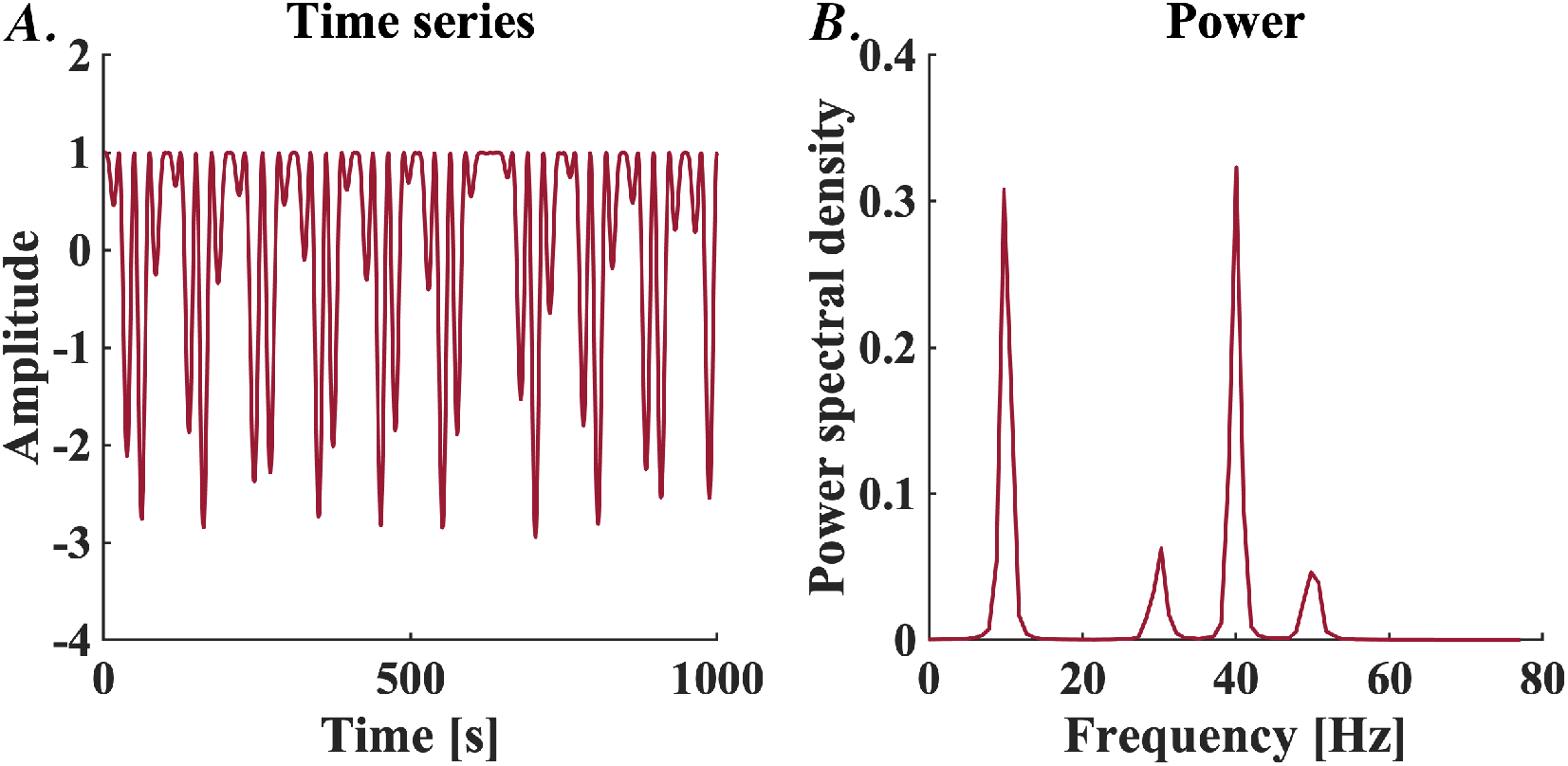
Example epochs of the simulated data. A) Example epoch of the simulated time series, and B) the corresponding power spectral density spectrum of the simulated signal.

Fig. 3 reveals the core of our argumentation why the filter bandwidth of the amplitude component should not be twice as large as the phase component. Fig. 3A depicts PAC with a fixed frequency of the phase component (10 Hz) and an amplitude component with a varying centre frequency and width. No PAC is observed for a bandwidth below 10 Hz. At a bandwidth of 10 Hz, two peaks around 35 and 45 Hz appear. Increasing the bandwidth further leads to smearing of both peaks and eventually to a merge of the two peaks at a bandwidth of (roughly) 20 Hz. In fact, the mere reason why PAC at 40 Hz can be observed is because the peaks at 35 and 45 Hz merge together.

**Figure 3:**
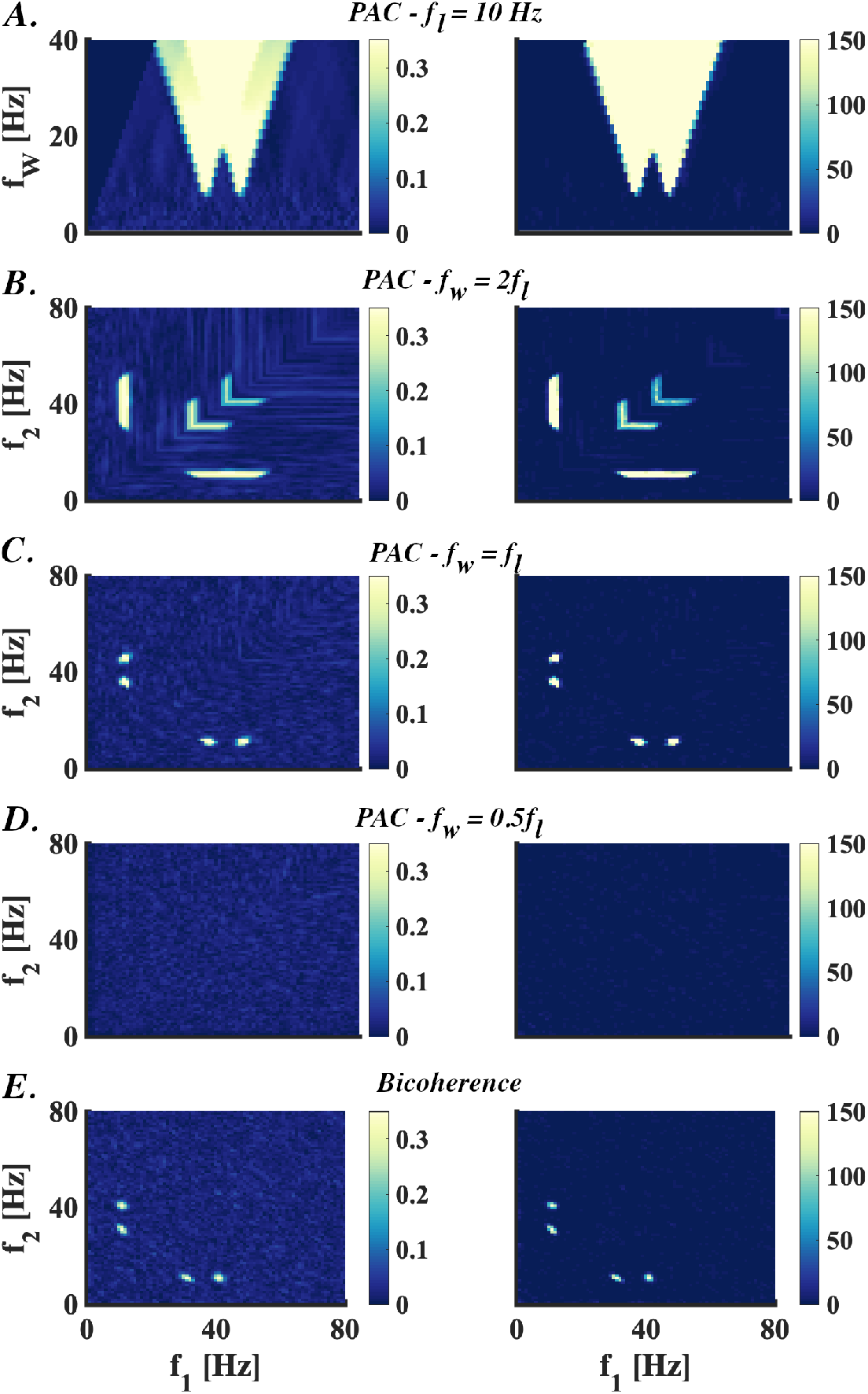
Phase-amplitude coupling and bicoherence in the simulated data. Left-hand plots represent the mean and right-hand plots the ratios between the actual cross-frequency coupling and surrogate coupling. A) Phase-amplitude-coupling (PAC) as a function of the amplitude component and its filter bandwidth. Here, the frequency of the phase component was fixed at 10 Hz. This representation reveals that two peaks of PAC can be observed at a bandwidth of 10 Hz while a bandwidth of 20 Hz reveals a smeared version. B-D) PAC as a function of the phase and amplitude component choosing a bandwidth that is double as large as (2:1-ratio), equals (1:1-ratio) or is half as large as (0.5:1-ratio) the frequency of the phase component. The representations reveal a clear dependency of PAC on the bandwidth of its amplitude component. E) Bicoherence representation yielding significant cross-frequency coupling at 30 and 40 Hz. Cross-frequency coupling in bicoherence matches the PAC estimate comprising a bandwidth that equals the phase component frequency. PAC: phase-amplitude-coupling; *f_W_*: filter width; *f*_1_: low frequency; *f*_2_: high frequency.

This becomes particularly evident when presenting PAC as a function of the frequencies of its phase and amplitude component. When the amplitude and phase frequency are in a ratio of 2:1, a smeared PAC can be observed ranging from 30-50 Hz with a clear peak at 40 Hz (Fig. 3B). If we, however, select a bandwidth with a 1:1-ratio then PAC reveals the two peaks around 40 Hz while the 40-Hz peak disappeared (Fig. 3C). PAC vanishes if the bandwidth becomes substantially smaller than the low frequency, which is shown here for a 0:5-1-ratio (Fig. 3D). In other words, the previously observed PAC at 40 Hz and the neighbouring peaks at 35 and 45 Hz disappears. The latter emphasizes the pivotal role of including at least one of the side frequencies at 30 and 50 Hz in the estimation of PAC. In the view of bicoherence, the observation of phase-amplitude coupling would be rather different compared to PAC (Fig. 3E). The bispectrum is the third order moment of the spectrum and measures the coupling between signals at three frequencies. Given the simulated data, two things could happen in terms of observing coupling between frequencies. There will be coupling between either 10, 30, and 40 Hz or 10, 40, and 50 Hz. In the bispectral analysis, bicoherence reveals most comparable results to the PAC estimated with a 1:1-ratio, except for a frequency shift of 5 Hz (30 and 40 Hz for bicoherence versus 35 and 45 Hz for PAC). This frequency shift is due to a different convention for the meaning of the frequencies. For example, bicoherence at a frequency pair at 10 and 30 Hz denotes coupling between 10, 30 and 40 Hz. The corresponding signal can be observed with PAC with 1:1 ratio with the filter for the high frequency part centred at 35 Hz such that the lower and upper end of that filter, having a width of 10 Hz, include the rhythms at 30 and 40 Hz.

### 3.2. Empirical electroencephalographic data

The human EEG data yield coupling at the alpha-band and its higher harmonics as revealed by bicoherence and PAC (Fig. 4). Like the simulated data, the PAC estimated on the EEG data highlight clear dependency on the bandwidth of the amplitude component (Fig. 4A). When the frequency of the phase component is constrained to be constant (here, set at 10 Hz), two peaks appear at a bandwidth of the amplitude component that is close to 10 Hz. There is hardly any (significant) PAC below a bandwidth of 10 Hz. A bandwidth enlargement above 10 Hz leads to an increased spectral window in which we can observe significant PAC and the two coupling peaks get smeared out over a broader frequency range. This ultimately results in the merging of the two peaks around a bandwidth of 15 Hz. Note that the bandwidth of the amplitude only is 1.5 times as large as the phase component frequency. Such cross-frequency coupling patterns also emerge when representing PAC over its phase and amplitude component while having the bandwidth depending on the phase component. The coupling is rather smeared and broad (i.e. roughly between 10 and 25 Hz) when the spectral representations are computed with a bandwidth of twice the frequency of the the phase component, the 2:1-ratio (Fig. 4B). On the contrary, multiple (disentangled) PAC peaks emerge when lowering the bandwidth to a 1:1-ratio relative to the frequency of the phase component (Fig. 4C). A focal PAC peak can be observed around 18 Hz with higher harmonics around 28 Hz. This emphasizes the essential role of the filter bandwidth in a 1:1-ratio when one is interested in phase-amplitude coupling in a small frequency band. The bandwidth size shrinkage leads to a PAC-modulation from broad-band smearing across a frequency band to multiple narrowed peaks. If the bandwidth is narrowed even more to a 0.5:1-ratio, no PAC can be detected anymore (Fig. 4D). Bicoherence spectrally identifies significant phase-amplitude coupling at the alpha rhythm (11 Hz) (Fig. 4E). Moreover, higher harmonics at 22 Hz are found as well. As expected, the 11-Hz peak is much more prominent than the higher harmonics. Compared to the PAC-estimates, bicoherence spectrally localised separated peaks, which were evident in the PAC-estimates with a 1:1-ratio. Here again, bicoherence is therefore most comparable to the PAC-estimates with a filter bandwidth that equals the low frequency component.

**Figure 4:**
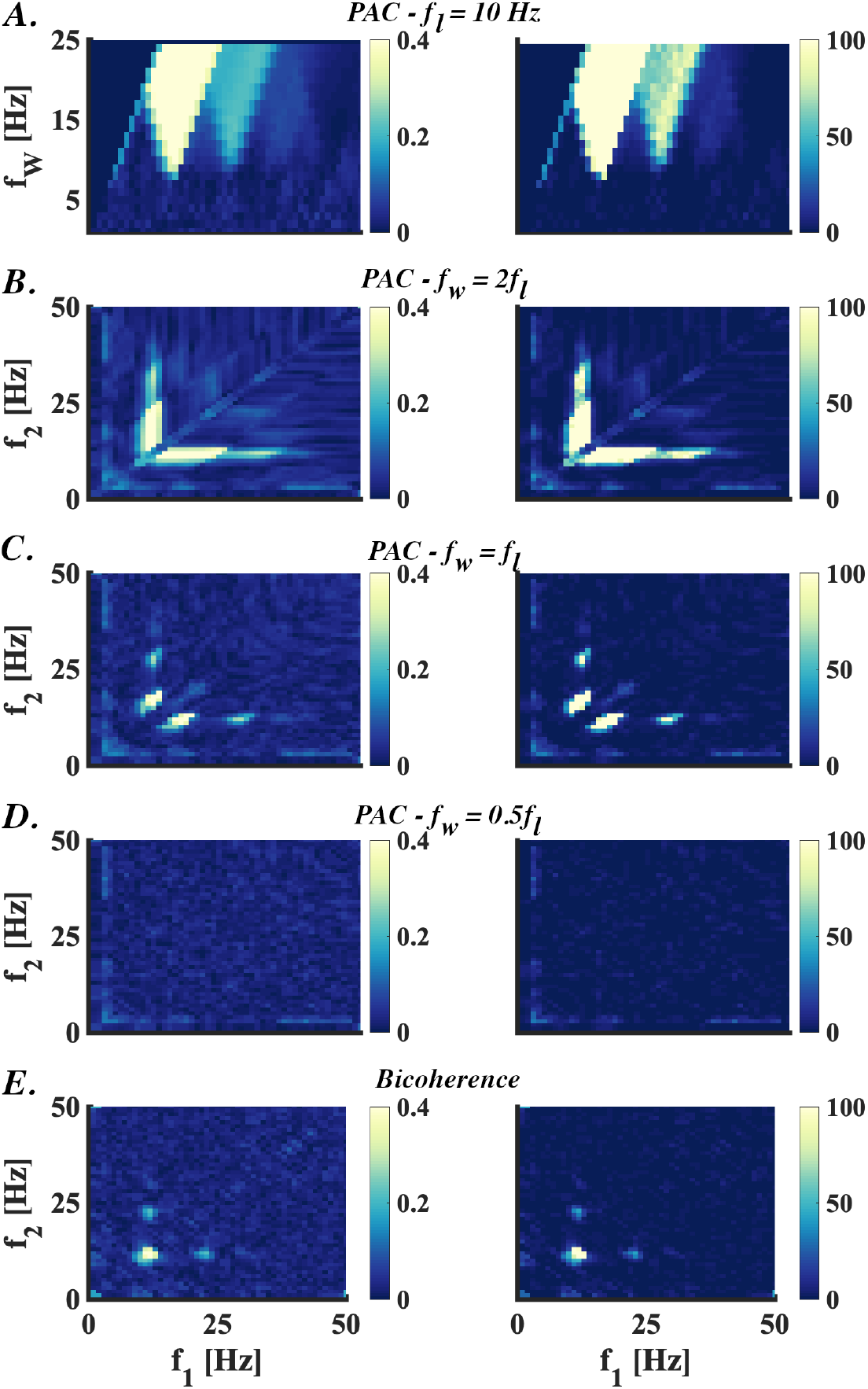
Phase-amplitude coupling and bicoherence in the human electro-encephalographic data. Illustrative coupling estimates examples of the electro-encephalographic data of single participant (channel 17 of subject 8). Left-hand plots represent the mean and right-hand plots the ratio values using the surrogate data. A) Phase-amplitude coupling (PAC) as a function of the amplitude component and its bandwidth. The low-frequency component was fixed at 10 Hz as this frequency corresponded to the power peak in the alpha range. B-D) The PAC representations based on a broader (2:1-ratio), equalling (1:1-ratio), or narrowed (0.5:1-ratio) bandwidth underlines the importance of using filter settings in a 1:1-ratio. Here, the broad-band smearing results in merging multiple peaks. E) Bicoherence spectrally localise the peaks 11 and 22 Hz like the PAC-estimates with a narrowed bandwidth. PAC: phase-amplitude coupling; *f_W_*: filter width; *f*_1_: low frequency; *f*_2_: high frequency.

### 3.3. Empirical local field potential data

The LFP data yield theta-gamma coupling resembled between 4-8 and 40-100 Hz. Like the simulation and EEG findings, coupling smearing across frequencies manifests in PAC when increasing the bandwidth frequency and fixating the phase component frequency (Fig. 5A). While the coupling at higher frequencies (>40 Hz) is hardly affected by the bandwidth, broadband smearing of the two low-frequency coupling peaks starts at frequencies of 6-7 Hz. Note that this frequency is (nearly) equivalent to the phase component frequency. As the coupling in the EEG data also highlighted, the merging of peaks appears at a bandwidth that is lower than twice the frequency of the phase component. Eventually, the low- and high-frequency coupling converges into one broad frequency range of coupling when the bandwidth is increased even further, for example, up to four times the phase component.

**Figure 5:**
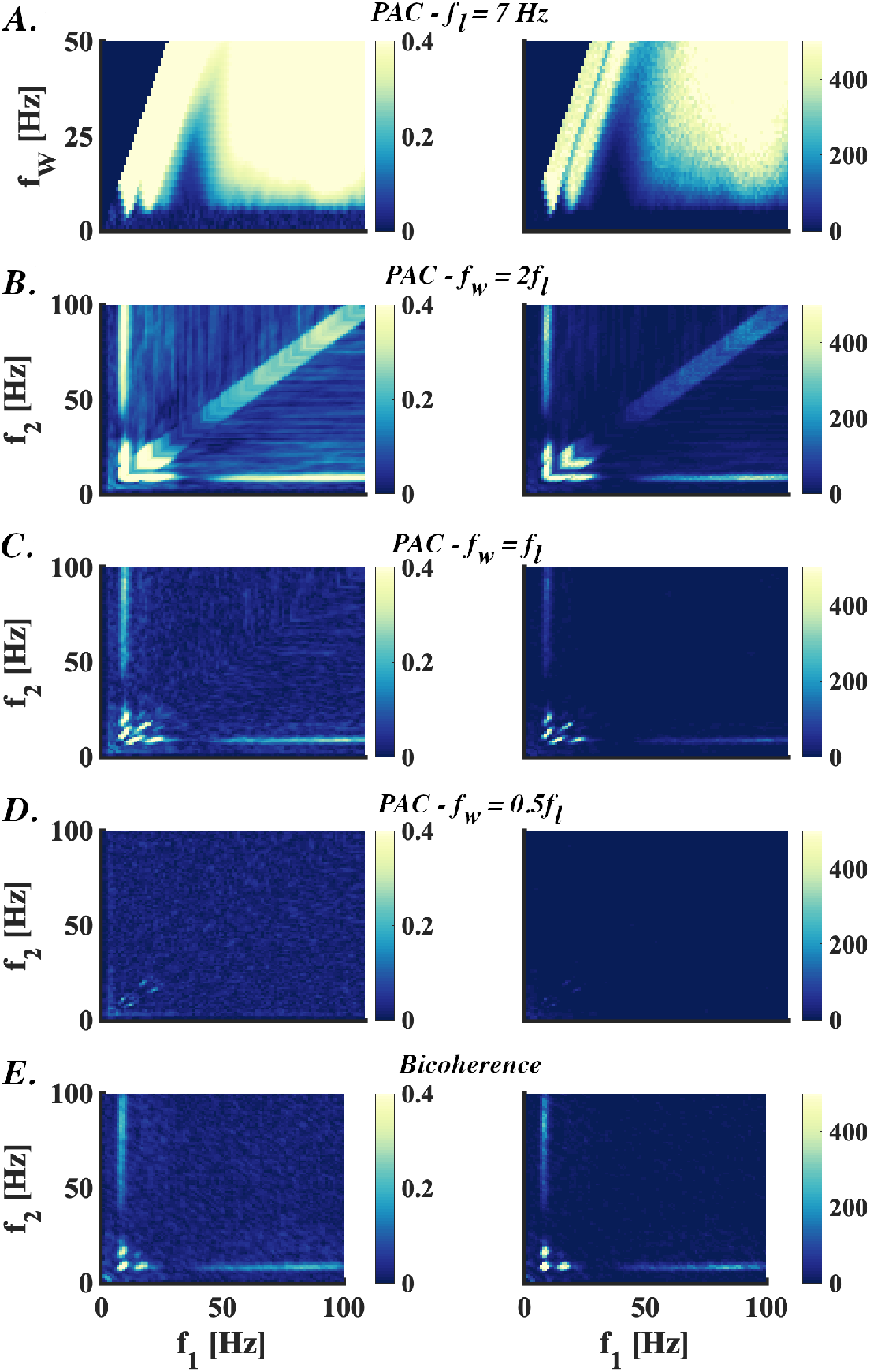
Phase-amplitude coupling and bicoherence in the local field potentials data of rats. Illustrative examples of the coupling estimates of the local field potentials data of a single rat (channel 16 of rat 7). Left-handed plots represent the mean coupling. Right-handed representations show the statistics of the surrogate data (here, the ratios between the coupling and surrogate coupling). A) Phase-amplitude coupling (PAC) that depends on (the bandwidth of) the high-frequency component while the frequency of the low-frequency component is fixed. To do so, the bandwidth was fixed at 7 Hz (i.e. the frequency where most coupling was expected). B-D) When the ratio between the phase component frequency and bandwidth of the amplitude component is set at a 2:1-, 1:1-, or 0.5:1-ratio, the (low-frequency) coupling reveals either smeared, non-smeared or non-existent PAC. The 1:1-ratio PAC establishes two low-frequency coupling peaks. E) These cross-frequency coupling peaks are also accurately identified by bicoherence. PAC: phase-amplitude-coupling; *f_W_*: filter width; *f*_1_: low frequency; *f*_2_: high frequency.

An amplitude bandwidth with a ratio of 2:1 to the phase component frequency results in broadly-smeared low-frequency coupling from 10 to 25 Hz (Fig. 5B). If we take a bandwidth that matches the phase component (i.e. the 1:1-ratio), peaks around 13 and 20 Hz can be recognized (Fig. 5C). Likewise, bicoherence is able to localise these two low-frequency coupling peaks (Fig. 5E). These peaks are around 9 and 16 Hz. For both PAC and bicoherence, and as displayed in Fig. 5A, the high-frequency coupling between 40 and 100 Hz does not depend on the bandwidth frequency and remains unchanged. This component only vanishes when PAC is estimated with 0.5:1 ratio for the bandwidth of the amplitude component (Fig. 5D).

## 4. Conclusion

Much of the brain’s dynamics is related to nonlinear neural activity. Cross-frequency coupling measures are sensitive to identify such neural processes. Here, we evaluated the phenomenon of phase-amplitude coupling from a bispectral point of view. Our main theoretical finding is that PAC and bispectra are not only related but are literally equivalent for an appropriate choice of the filter to estimate the high frequency amplitude, and even then PAC is just a special case of bicoherence. As a mathematical result, this finding is an idealization, and there can still be rather minor deviations in practice for several reasons which we would like to summarize:

1. We have shown the equivalence for PAC and bispectra for infinite frequency resolution, but frequency resolutions are finite in practice with details depending on the implementations.
2. Equivalence is shown when PAC can be expressed as a third order statistical moment, which requires that the low frequency phase is weighted by the low frequency amplitude and that the coupling to the squared amplitude is studied rather than the amplitude itself. Whether such requirements are fulfilled depends on the version of PAC.
3. Mathematical relations refer only to the numerators of the coupling measures, i.e. the bispectra. However, we found in empirical studies that the normalizations do not affect the principal finding.

Our own implementation of PAC, i.e. coherence between signal and high frequency amplitude at some low frequency, deviated from the above idealization with regard to all three aspects. We still found that results for PAC and bicoherence are always very similar using a proper choice of filters for the calculation of PAC.

From these findings we can derive some recommendations how to choose filters for the calculation of PAC. When cross-frequency phase-amplitude interactions are estimated with PAC, filter settings substantially affect the coupling. Although it is already well established that the use of adequate filter bandwidths is critical to estimate PAC accurately (Aru et al. [2015]; Berman et al. [2012]), there is no consensus in terms of the optimal filter settings. A recent study recommended to have the bandwidth of the amplitude component at least double as large as the low-frequency component (Aru et al. [2015]). Here, we proved that this recommendation is not trivial from a bispectral perspective. In particular, such filter settings hamper coupling estimation when one is interested in neural dynamics within a narrowed frequency range. As revealed by both simulation and empirical findings, cross-frequency phase-amplitude coupling is evident in both PAC and bicoherence estimates. However, as expected, the PAC dynamics were clearly modulated as a function of the filter settings (i.e. the bandwidth of its amplitude component). Specifically, we show that the frequency range over which significant PAC can be observed decreases when the bandwidth of the high-frequency component decreases. PAC-estimations comprising a bandwidth that is twice as large as the low-frequency component results in smearing of the phase-amplitude coupling over a broad frequency range. In other words, the spectral resolution of PAC substantially decreases when taking a bandwidth of the amplitude component that is too large. It then becomes difficult to distinguish wide band phenomena from higher harmonics. Bicoherence and PAC provide comparable results in case the bandwidth of the amplitude component is narrowed to match the frequency of the phase component. When the bandwidth of the amplitude component decreases even further, the likelihood that carrier frequencies are not included increases, meaning no PAC can be observed at all.

A further advantage of bicoherence over PAC is that the filtering of the phase and amplitude component at many isolated frequencies and frequency bands makes the estimation of PAC computationally heavy. In particular, the computation of a sufficient number of surrogate data during statistical assessments is time-demanding. In contrast, bicoherence is based on the fast Fourier transforms at three isolated frequencies. The latter, therefore, has the advantage that there is no necessity to filter signals before connectivity estimation. We conclude that, although PAC and bicoherence are measures to estimate cross-frequency phase-amplitude coupling, bicoherence has the advantage that it does not have to be estimated with additional pre-processing steps. Likewise, the filter settings of PAC’s amplitude component bandwidth should be chosen with precaution and should be equal to the frequency of the low-frequency (i.e. the phase) component. This is especially true for neural rhythms operating in narrow frequency bands.

## Acknowledgements

CSZ was financially supported by the ERC-Starting Grant “Learn2Walk” (ERC-Stg No. 715945) and NWO (Netherlands Organisation for Scientific Research) VIDI-grant “FirSTeps” (No. 016.156.346), both awarded to dr. Nadia Dominici. CSZ would like to thank the Faculty of Behavioural and Movement Sciences (Vrije Universiteit Amsterdam) for granting him a travel fund to visit the University Medical Centre Hamburg-Eppendorf. This research was also partially funded by the German Research Foundation (DFG, SFB936/Z3 and TRR169/C1/B4).

